# Biodiversity and distribution of sea anemones (Cnidaria, Anthozoa, Actiniaria) in Peru

**DOI:** 10.1101/2022.08.25.505348

**Authors:** Allison Durand, Deysi Valdivia-Chávez, Víctor Aramayo

## Abstract

Diverse and abundant sea anemones are common in shallow marine areas. Detailed biodiversity analysis in Peru are comparatively scarce. To contribute to the biodiversity inventory and distribution information of this taxa, we analyzed exhaustively the available bibliography in Peruvian waters. A total of 23 anemone species were identified, distributed in 68 localities, and grouped into 1 Order (Actiniaria), 6 Families (Actiniidae, Actinostolidae, Aiptasiidae, Isanthidae, Phelliidae and Sagartiidae) and 20 Genera. The most reported species are *Anthothoe chilensis* (37 references), *Phymactis clematis* (28), *Phymanthea pluvia* (27), *Oulactis concinnata* (18), and *Antholoba achates* (15). Lima is the region with the highest number of publications, followed by La Libertad, Piura, Lambayeque, and Ancash. *Anthothoe chilensis* occurs in almost all the Peruvian coastal regions. On the contrary, *O. concinnata* has been primarily observed in Lima, while *A. achates* occurred only in the southern regions (Ica, Arequipa, and Moquegua). Rocky substrates (~55% records) seem to be the most suitable habitat for sea anemones in Peru, corresponding to exposed (e.g. vertical walls) and sheltered zones (e.g. rocky crevices, caves, under rock areas). Although most of the species in Peru exhibit a relatively wide spatial distribution, our results suggest that there are several regions with little or no research efforts. Despite a growing study effort over the past 30 years (>50% of biodiversity reported), the current biodiversity status for this group is still unclear. A significant effort is needed to better analyze occurrence patterns and unveil new species regarding a changing environmental scenario with human influence.

## INTRODUCTION

The current taxonomic classification of Actiniaria (Rodríguez *et al*., 2014) includes two suborders: Anenthemonae and Enthemonae. These suborders exhibit anatomical differences. In the first suborder, a unique arrangement of mesenteries is characteristic; while the second suborder possesses a typical body scheme, mostly hexamerous cycles, in which pairs of mesenteries arise in the exocoels (Rodríguez *et al*., 2014). To the untrained eye, actiniarians look very similar; however, the so-called true anemones account for approximately 1200 species grouped in 46 families and are represented within the subclass Hexacorallia, class Anthozoa (Brusca, 1980; Daly *et al*., 2007).

Sea anemones constitute a conspicuous and diverse group in marine environments across all depths and latitudes (Daly *et al*., 2007; Fautin, 2013; Rodríguez *et al*., 2014). These organisms survive mostly adhered to hard substrates, but several species exhibit both a commensalism strategy (e.g. epibiosis on hermit crabs) (Ruppert & Barnes, 1996), and burial behaviour (sand and/or mud) (Häussermann & Försterra, 2009). A large number of polypoid-shaped species grow in colonial or solitary stages of surviving, yet clonal aggregations stand out as a notoriously efficient and successful strategy for rapid growth, space competition, and dispersal of individuals (Fautin, 2013).

Physical factors modulating the distribution and aggregation patterns of sea anemones are substrate availability and local oceanographic conditions (e.g. transparency, turbulence, and current patterns). Nevertheless, intra- and interspecific competition for food and space have largely been known as equally crucial for these organisms (Chintiroglou & Koukouras, 1992). Early-life survival and ecological connectivity also play a critical role in the success of the settlement of sea anemones species and their distribution among marine regions (Ocaña *et al*., 2007; Watson *et al*., 2018).

Because of their high metabolic sensitivity along their life cycle, some species have been proposed as key bioindicators of pervasive changes in the chemistry of surrounding waters and the overall marine environment health (Linton & Warner, 2003; Duckworth *et al*., 2017).

Although frequently observed living at rocky and sandy shores substrates, exhibiting both patchy and highly dense distributions along depth gradients, in-deep studies addressing the sea anemone biodiversity and ecology remain scarce in Peruvian waters. Most of the available information comes from sporadic reports for specific sites as a result of compilatory efforts (e.g. Novoa *et al*., 2010; Hooker *et al*., 2011) or general benthic inventories (e.g. Paredes *et al*., 1999; Uribe *et al*., 2013). There is no specific research aiming the sea anemones as a target group in Peru. As such, specific information for these organisms is quite scarce and scattered, limiting our opportunities to assess the ecological relevance of these populations and analyse their current status. To contribute to the study of this benthic group, in this paper we perform an exhaustive bibliographic review to document the biodiversity and distribution of sea anemone species reported in Peru.

## MATERIAL AND METHODS

### Study area

Information and data published on sea anemones from 68 localities in 10 Peruvian coastal regions were used, the distribution of the most representatives is shown in Figure 1. This spatial scale covers a study area comprising practically the entire Peruvian coast, including both cold and tropical marine ecosystems of this region, and different types of coastal habitats (beaches, islets, islands). The Peruvian waters are recurrently affected by interannual events such as El Niño (EN) and, from shallow to deeper areas, by an intense oxygen minimum zone (OMZ, dissolved oxygen < 0.5 ml.l^−1^) that modulates the abundance and development of both pelagic and benthic communities (Tarazona *et al*., 2003; Bertrand *et al*., 2011).

**Figure 1.**
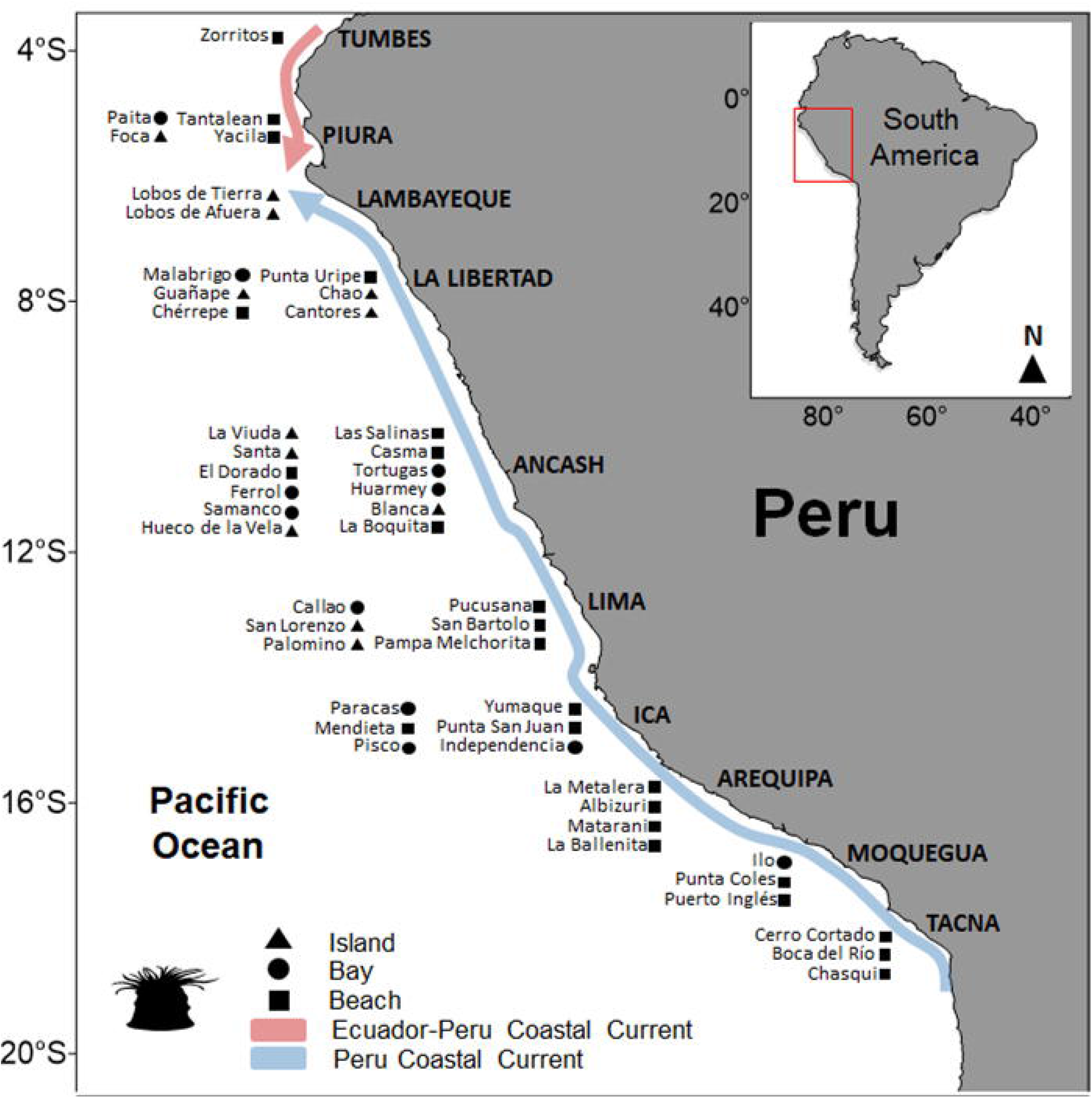
Study area and most representative localities of sea anemones in Peru. Symbols accompanying each locality describe the type of site (island, bay, or beach) in which sea anemones have been reported. Though indicated as an island, Cantores and Hueco de la Vela are inlets.

### Data sources and criteria employed

An exhaustive literature review was performed to list the sea anemone species reported at different benthic habitats in Peruvian waters. We focused on the true anemones only. The Ceriantharia and Hexacorallia subclasses (‘tube anemones’ and ‘colonial anemones’, respectively) were excluded since these groups are critically less documented in Peru.

Our geographic information range included data from both the equatorially influenced region in Northern Peru (approximately from the Peruvian northern border to ~5°S) and the cold region from Central and Southern Peru (Chaigneau *et al*., 2013), also named the Humboldtian region off Peru. Overall, we reviewed 110 scientific references that analyse several topics of sea anemones, including peer-reviewed papers, official (governmental) reports, MSc and/or PhD theses (when strictly necessary), and even seminal reports (e.g. Verrill, 1867). Grey literature was explicitly excluded in this work (conference abstracts, nonreferenced technical reports, unsourced or poorly documented online material, etc.) Authoritative online resources/databases were used to cross/confirm local references in some cases and verify the global incidence of some species; for instance, the World Register of Marine Species (WoRMS: http://www.marinespecies.org/), the Ocean Biodiversity Information System (OBIS: https://obis.org/), the Global Biodiversity Information Facility (GBIF: https://www.gbif.org/) and the Biodiversity Heritage Library database (BHL: https://www.biodiversitylibrary.org/).

When the geographic coordinates range and/or specific locality of sampling was not indicated, we used the local name of the city reported in the document as the main reference. Our list includes the region name followed by the specific locality (in parentheses). Due to missing information regarding the bathymetric range of some species, our discussion of the bathymetric distribution was based only on reliably known information. Finally, to avoid confusion when indicating scientific names, after its first mention, we decided to use the extended name of *Anthothoe chilensis* to differentiate it from *Actinostola chilensis*.

## RESULTS

### Sea anemone biodiversity and latitudinal distribution

A total of 23 species have been reported for the Peruvian coast, grouped into 1 order (Actiniaria), 1 superfamily with a temporary name (Actiniaria *incertae sedis*), 6 families (Actiniidae, Actinostolidae, Aiptasiidae, Isanthidae, Phelliidae and Sagartiidae) and 20 genera (Figure 2A, Table 1). The most diverse families were Actiniidae (11 spp., 50% of total), Actinostolidae, and Sagartiidae (both with 3 spp., 13.6%). Both *Anactis picta* (Lesson, 1830) and *Paractis peruviana* (Lesson, 1830) are not currently assigned to a specific family; instead, they have been temporally placed under the category Actiniaria *incertae sedis* (Table 1). These last two species have been mentioned in early bibliographic references only (Verrill, 1867; Pax, 1912). Contemporaneously, Fautin (2013, 2016) cited the same authors when referring to these species.

**Fig. 2.**
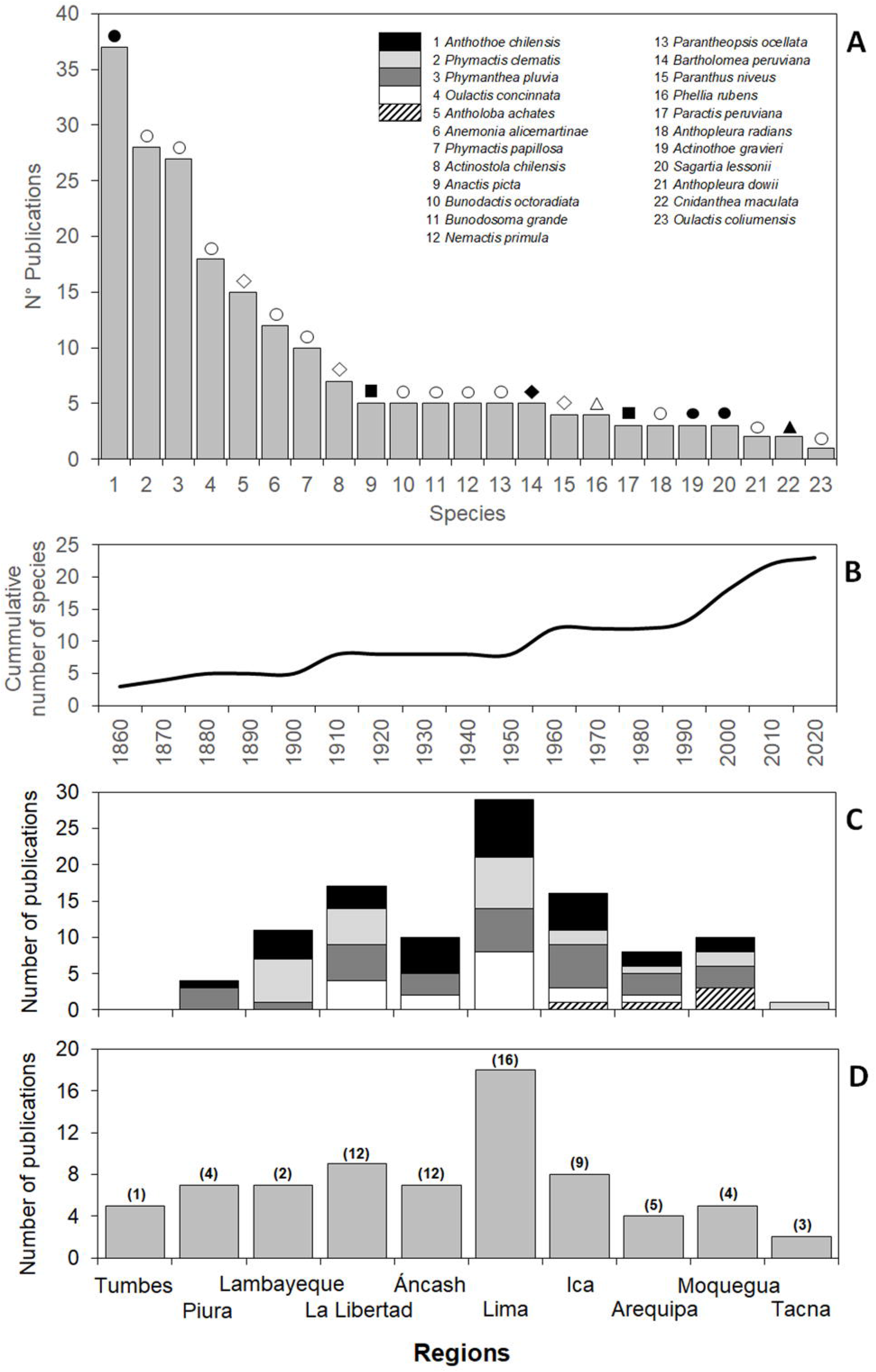
A) Number of publications by species of sea anemones reported in Peru. Symbols indicate the families Sagartiidae (black circle), Actiniidae (white circle), Actinostolidae (white rhombus), Aiptasiidae (black rhombus), Actiniaria *incertae sedis* (black square), Phelliidae (white triangle) and Isanthidae (black triangle). B) The cumulative number of species over time from 1860 to 2020. Regional variability of C) The number of publications for the top five most studied sea anemones species, and D) The sampling effort (No. publications) along the Peruvian coast, number of localities between parenthesis.

**Table 1.**
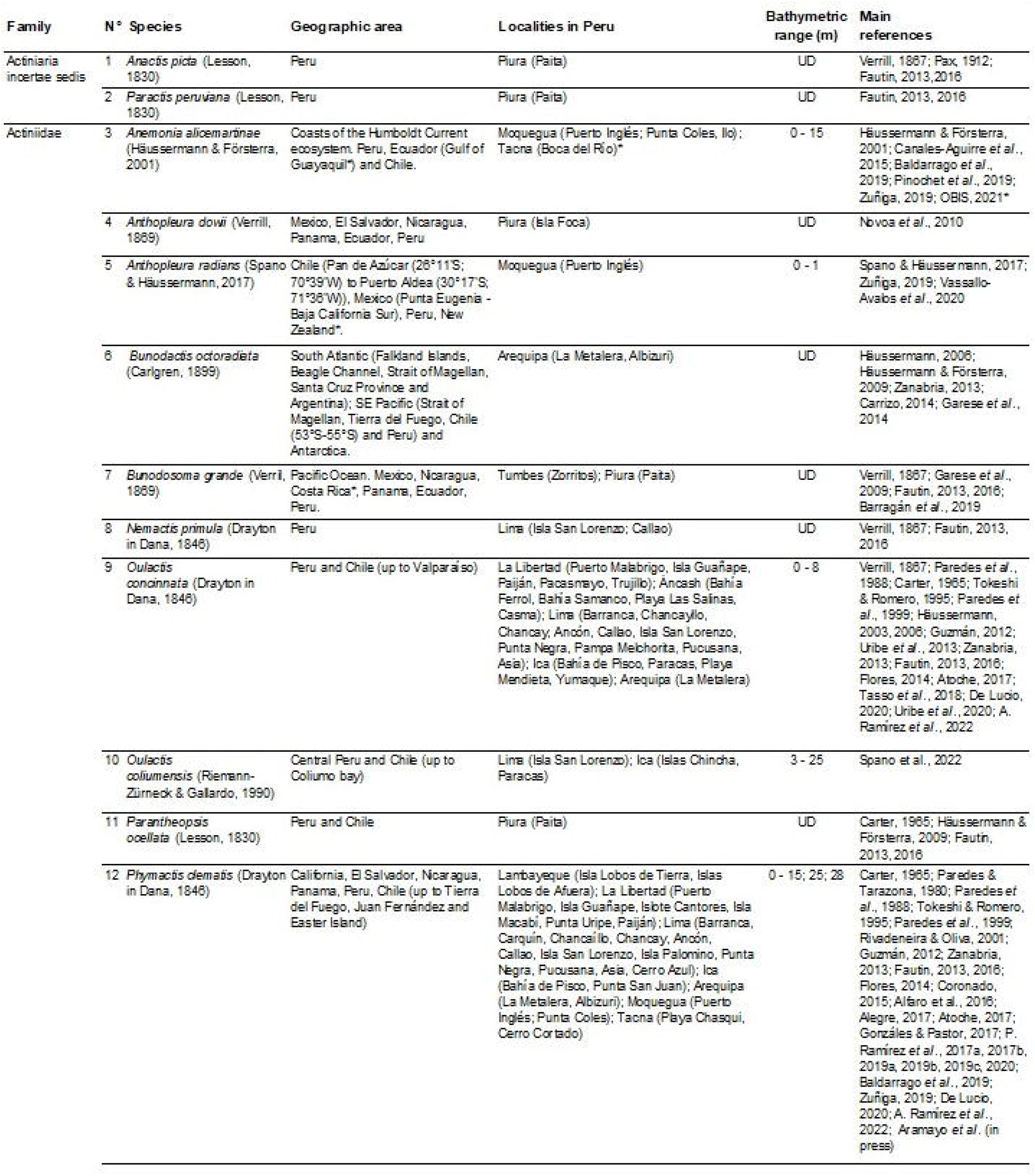
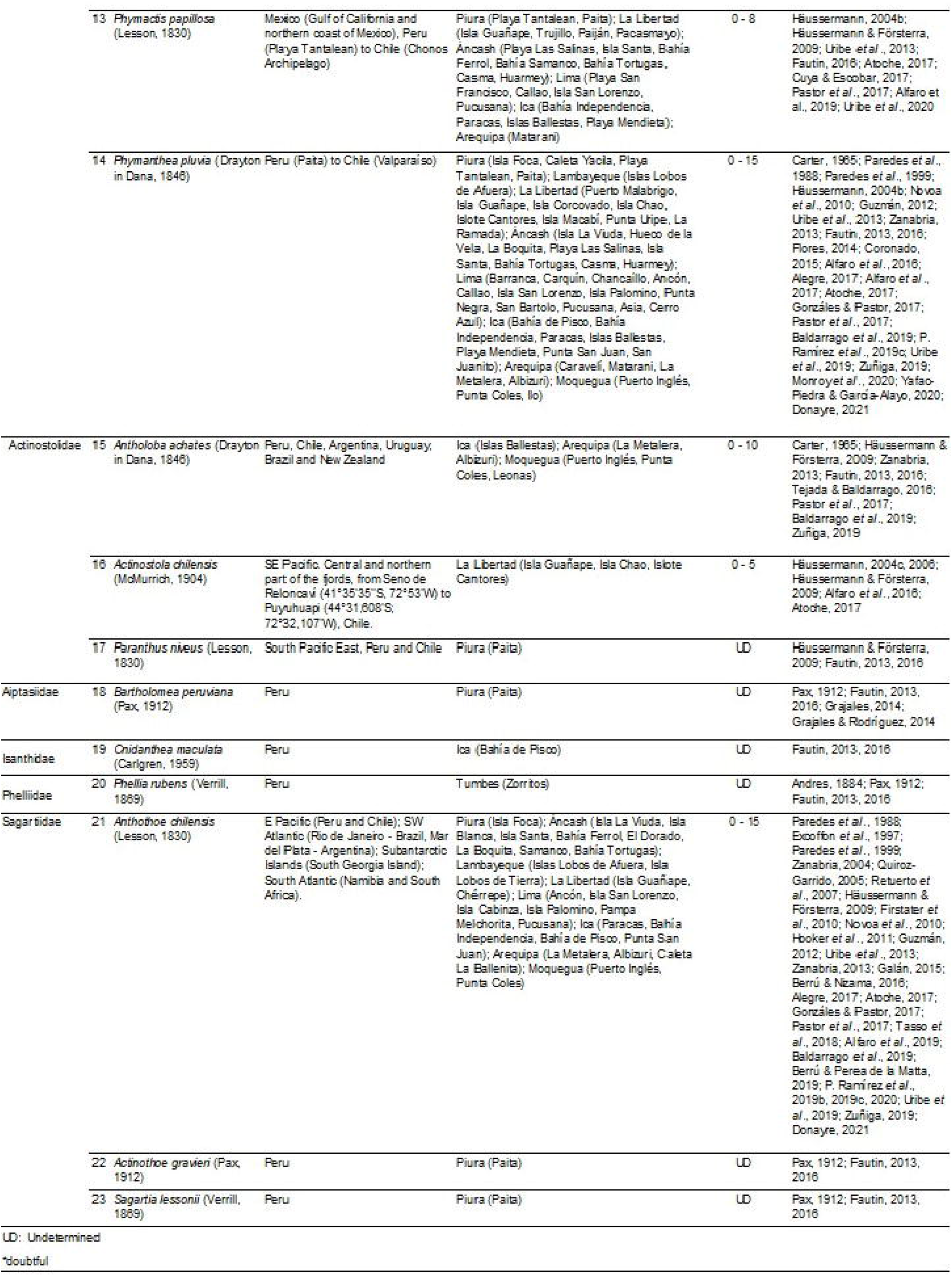
Sea anemones species from Peru, considering taxonomic information, geographic distribution, the total number of localities, and bathymetric range. UD: undetermined. Symbol (*) indicates doubtful information.

Out of 23 species identified, the top five exhaustively reported are *Anthothoe chilensis* (Lesson, 1830) (37 references), *Phymactis clematis* (Drayton in Dana, 1846) (28), *Phymanthea pluvia* (Drayton in Dana, 1846) (27), *Oulactis concinnata* (Drayton in Dana, 1846) (18), and *Antholoba achates* (Drayton in Dana, 1846) (15) (Figure 2A). Major contributions to the scientific references of these and other species have been authored by Häussermann & Försterra (2009) and Fautin (2013, 2016). Moreover, our time series for reported species in Peruvian waters (1860-2020) revealed very low progress, with long periods of research inactivity in terms of new species discoveries or new records, with slight changes every 50 years approximately, and a clear shift over the last 30 years only (Figure 2B).

Our findings indicate that the top five reported species (~22%) have a broad spatial distribution (Figure 2C, Table 1). For instance, studies concerning *Anthothoe chilensis*, *Phymactis papillosa* (Lesson, 1830), *O. concinnata*, *P. clematis*, and *P. pluvia* reveal the extensive distribution of these species, especially in intertidal zones throughout the Peruvian coast. Conversely, about 50% of identified species for Peru, namely *Anactis picta* (Lesson, 1830), *Paractis peruviana* (Lesson, 1830), *Anthopleura dowii* (Verrill, 1869), *Parantheopsis ocellata* (Lesson, 1830), *Actinostola chilensis* (McMurrich, 1904), *Paranthus niveus* (Lesson, 1830), *Bartholomea peruviana* (Pax, 1912), *Phellia rubens* (Verrill, 1869), *Actinothoe gravieri* (Pax, 1912), *Sagartia lessonii* (Verrill, 1869), and *Bunodosoma grande* (Verrill, 1869) exhibit a latitudinal distribution restricted to the northern regions. Less-studied species such as *Anemonia alicemartinae* (Häussermann & Försterra, 2009), *Bunodactis octoradiata* (Carlgren, 1899), *Anthopleura radians* (Verrill, 1869), *A. achates* and *Cnidanthea maculate* (Carlgren, 1959) seem to be restricted to the southern coast (~23% of spp.) Likewise, *Nemactis primula* (Drayton in Dana, 1846) has been exclusively reported for the central region (Lima, ~5%). Recent reports (Spano *et al*., 2022) indicate that *Oulactis coliumensis* (Riemann-Zürneck & Gallardo, 1990) inhabit the central and southern coasts.

The highest number of scientific publications (18) corresponds to the Lima region, with more than twice the studies developed for other regions (Figure 2D). It is followed by La Libertad (9), and the regions of Piura, Lambayeque, and Ancash (7). However, the number of localities studied in each region is unevenly distributed, since some north-central regions (La Libertad, Ancash, and Lima) exhibit a high number of localities surveyed, compared to the southern regions such as Arequipa, Moquegua, and Tacna (Figure 2D). Consequently, the research effort (indicated by the number of publications) does not seem to have targeted those regions with a rather scarce number of studies.

### Habitats range

The description of habitat features (i.e. chemical and physical properties or the influence of gradients) is little analyzed, or it is absent, regarding sea anemones reports. Only 15 species (65%) have studies roughly offering habitat descriptions, while the other 8 (35%) have no specific data to discuss. Rocky substrates, like exposed (e.g. vertical walls) or sheltered zones (e.g. rock crevices, caves, intertidal pools), were the most frequently reported type of habitat (~55% records). Several species live indistinctively along the depth gradients inside the intertidal zone, whereas only a few permanently inhabit the subtidal shallow zone. It is worth noting the lack of bathymetric data for Peruvian waters, as we found information for 10 species only, from the Actiniidae, Actinostolidae, and Sagartiidae families (Table 1), inhabiting shallow environments (between 0 and 28 m). Notoriously, the vertical distribution of 13 species remains undescribed.

Several observations for the Peruvian regions of Ancash, Lima, Moquegua, and Ica suggest a shallow distribution for *Anthothoe chilensis* (Figure 3A), with a bathymetric range fluctuating from the intertidal down to 15 m depth (Table 1). Indeed, this species often inhabits the littoral and sublittoral zones forming small aggregations; tidepools, crevices, and exposed areas have been reported as its common shallow habitats. Even so, these organisms can be also found on mixed and biogenic substrates as well as living in kelp forests.

**Fig. 3.**
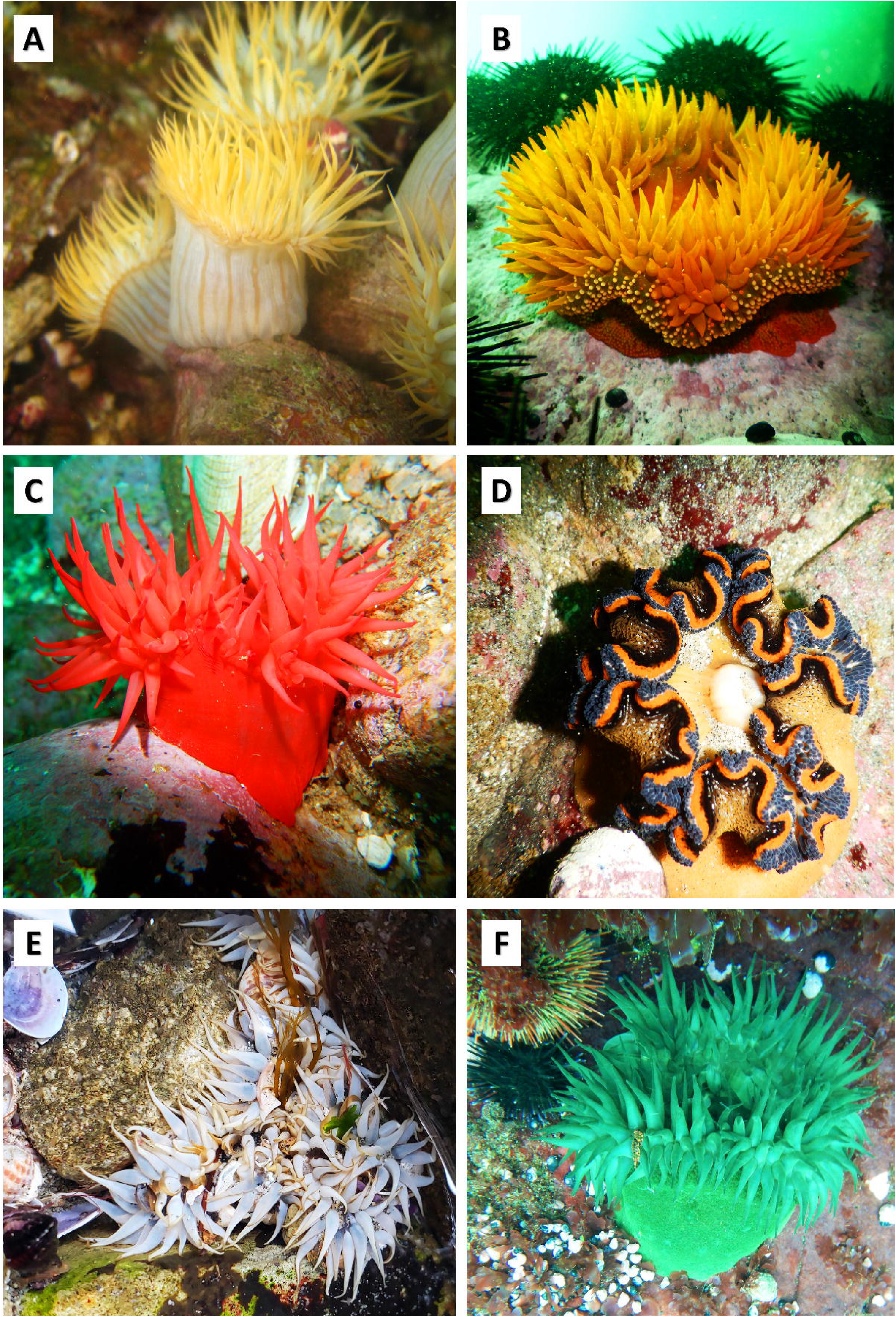
Representative sea anemones species inhabiting coastal areas in Peru. A. *Anthothoe chilensis*; B. *Phymanthea pluvia*; C. *Anemonia alicemartinae*; D. *Antholoba achates*; E. *Oulactis concinnata*; F. *Phymactis clematis*. Photo credits: A-D (R. Uribe), E (V. Aramayo), F (D. Baldarrago).

Overall, *P. pluvia* (Figure 3B) lives on rocky shores, shallow waters, and hard bottoms. Individuals are present in the lower intertidal and subtidal zone, in small caves or attached to vertical walls. Observations from the regions of La Libertad, Ancash, and Moquegua reveal that the bathymetric distribution of this species extends from the intertidal zone down to 15 m depth (Table 1); although also reported in other regions (Figure 2C), its vertical distribution is still poorly described. In the same sense, samples from the region of Moquegua indicate that individuals of *A. alicemartinae* (Figure 3C) live down to 15 m depth (Table 1), inhabiting both the intertidal and shallow subtidal zones, attached to bare rock surfaces, half-buried under the sand or even on floating macroalgae.

Despite being one of the most reported sea anemone species, *A. achates* (Figure 3D) remains insufficiently documented, with several local observations (D. Baldarrago, per. com.), and scarce registers of its occurrence derived from the southern coast only (Figure 2C). However, a study in the Moquegua region has reported its presence down to 10 m depth (Table 1). Other dwellers such as *O. concinnata* (Figure 3E) live among substrates like crevices or under the rocks of shallow hard bottoms from the intertidal to the shallow subtidal zones. Its occurrence in the Ancash region suggests a restricted vertical distribution ranging from the intertidal down to 8 m depth (Table 1). Unfortunately, despite having been reported in other regions, site-specific data is not available, and most information is addressed roughly indicating rocky habitats and exposed zones as representative coastal environments.

As for the bathymetric data available for *P. clematis* (Figure 3F), individuals have been observed down to shallow subtidal zones (15 m depth) in the Moquegua region; while specimens have been found in samples collected from sandy and muddy bottoms at 25 and 28 m depth in the Tacna region (See references in Table 1). Occurrences in other regions have also been registered but lack precise bathymetric data.

Furthermore, the unique bathymetric distribution data available for *A. chilensis* derives from samples obtained from vertical walls in Guañape Island (between 0 and 5 m depth) in the La Libertad region (Table 1). Likewise, *P. papillosa* has been reported throughout the Peruvian coast, but its bathymetric range (between 0 and 8 m depth) is known only for the Ancash region (Table 1). Finally, just a narrow bathymetric range was attributed for *A. radians* (0 – 1 m, Table 1) with occurrences on rocky shores of the Moquegua region.

The superfamily Actiniaria *incertae sedis* involves two species (*A. picta* and *P. peruviana*) with an unknown bathymetric range for Peru. Unfortunately, over the last two decades, there is no new data to resolve this uncertain status regarding the spatial distribution of these species and back up the original findings (see Fautin, 2013; 2016).

## DISCUSSION

### Species occurrence and biodiversity status

Despite 13 species shared by Peru and geographically near areas such as northern Chile (*A. alicemartinae, A. radians, B. octoradiata, O. concinnata, O. coliumensis, P. ocellata, P. clematis, P. papillosa, P. pluvia, A. achates, A. chilensis, P. niveus*, and *Anthothoe chilensis*), specific information for the Peruvian coast is comparatively scarce. Most of the data available derives from monitoring reports carried out only once in a particular locality (Hooker *et al*., 2011; Uribe *et al*., 2013; Gonzáles & Pastor, 2017; Pastor *et al*., 2017). Diversity and distribution research targeting sea anemones in Peru has not yet been conducted, in contrast to other efforts in the region (Lancellotti & Vasquez, 2000; Häussermann & Försterra, 2005; Häussermann, 2006). In fact, most occurrence reports, including taxonomy and ecological knowledge originate from overseas studies (e.g. Carter, 1965; Stotz, 1979; Excoffon *et al*., 1997; Häussermann & Försterra, 2001; Häussermann, 2003, 2006, 2004b, 2004c; Fautin, 2016; Spano & Häussermann, 2017; Pinochet *et al*., 2019). Nevertheless, our analysis suggests a gradual increase in interest in sea anemone biodiversity over time (Figure 2B).

Certainly, this is relatively recent, as representative species *P. clematis* and *Anthothoe chilensis* were reported for the first time in the 1980s (Paredes & Tarazona, 1980; Paredes *et al*., 1988). Since then, their occurrence in reports for the Peruvian coast has augmented considerably. Similar cases could follow a similar pattern; for instance, species such as *A. alicemartinae* reported for the first time in 2015 (Canales-Aguirre *et al*., 2015) and *O. coliumensis*, whose northward spatial extension was recently confirmed (Spano et al., 2022).

Species like *Anthothoe chilensis*, *P. clematis*, *P. pluvia*, and *O. concinnata* have been regularly mentioned in recent benthic diversity inventories and monitoring reports (Uribe *et al*., 2013; Alfaro *et al*., 2016, 2019; Gonzáles & Pastor, 2017; Pastor *et al*., 2017; Tasso *et al*., 2018; Baldarrago *et al*., 2019; Berrú & Perea de la Matta, 2019; Uribe *et al*., 2019; P. Ramírez *et al*., 2020, 2019b, 2019c, 2019a; A. Ramírez *et al*., 2022). Perhaps one explanation for their high occurrence in reports is the abundance of their populations in intertidal and shallow subtidal habitats. However, the lack of detailed registers in previous years providing information on this issue questions this explanation. Probably, specimens were only reported as “Cnidaria", “Actiniaria indet.”, “actiniarian”, “anemone", were not detected or simply confused with other species. This situation is common with local observations for some indeterminate, but frequently observed species that cohabit in the intertidal and shallow subtidal (~25 m). For instance, sea anemones still unidentified have been found in southern Peru attached to rhizoids or fronds of *Lessonia trabeculata* (Figure 4A-C) or even adhered to crevices in intertidal rocky shores (Figure 4D). These findings highlight the necessity of increasing taxonomic identification efforts to expand the actual insights of sea anemone’s diversity in southern regions.

**Fig. 4.**
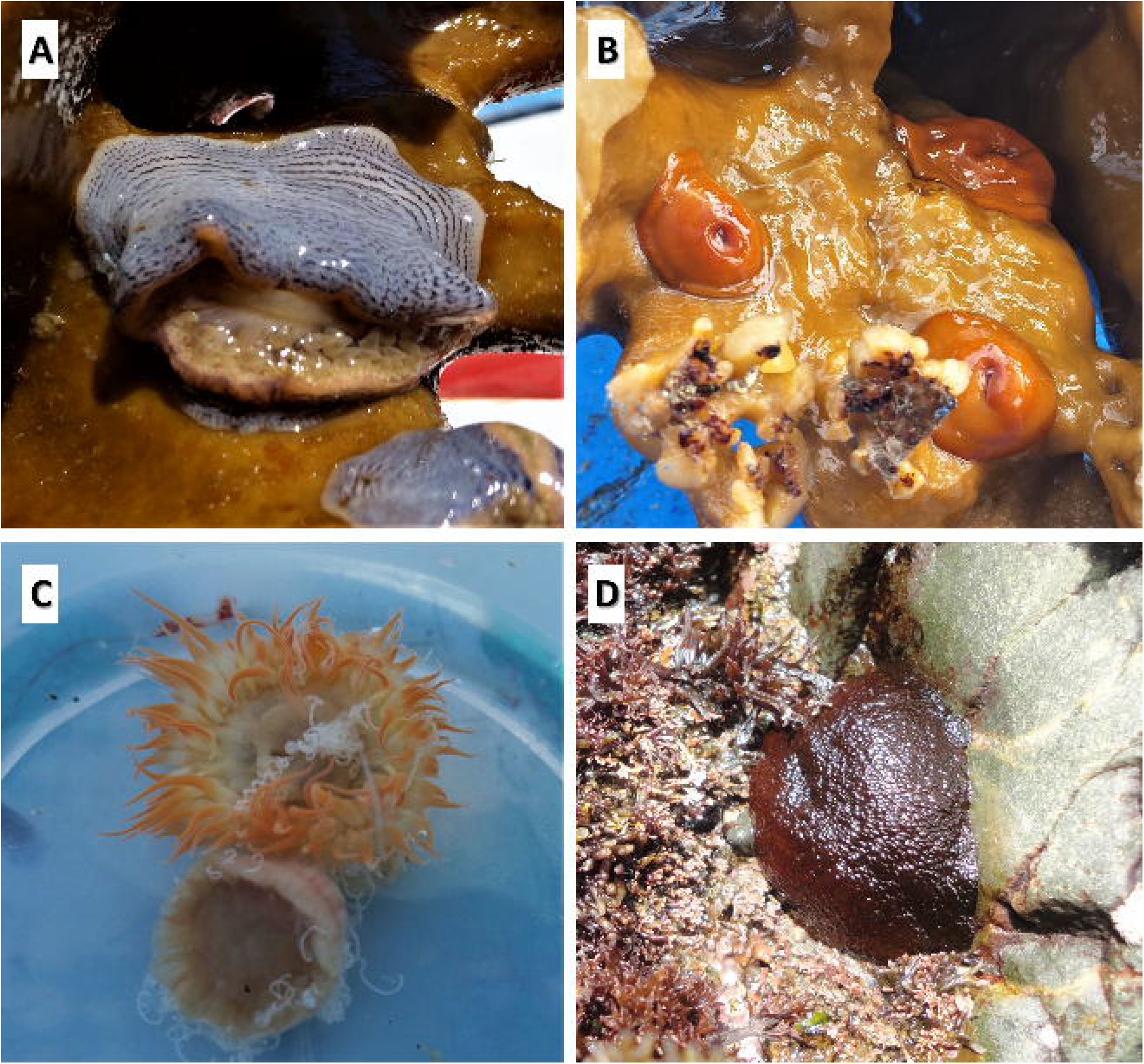
Unidentified species of sea anemones in southern Peru. Photo credits: A-D (D. Valdivia).

The restricted information on qualitative and especially quantitative data (e.g. bathymetry), is another bottleneck for dimensioning the distribution of sea anemone communities in shallow waters. Habitat data has been the most critical point hindering better descriptions, this is common in monitoring reports. For instance, punctual depth values of the occurrence of *P. clematis* (Table 1) in the subtidal were obtained as individuals were found casually in samples collected to study the soft-bottom macrobenthos in southern Peru (Aramayo *et al*., in press). Additionally, inaccurate bathymetric data has been found throughout our literature review. Among the scant information available, imprecise descriptions of the depth at which sea anemones have been found or sighted are unfortunately recurrent (e.g. Paredes & Tarazona, 1980; Paredes *et al*., 1999; A. Ramírez *et al*., 2019; De Lucio, 2020; P. Ramírez *et al*., 2019b, 2019c, 2019a). Thus, we possess a relatively high degree of uncertainty regarding the bathymetric distribution of several species, limiting our capacity to analyse the influence of depth on biodiversity.

As an example, sea anemone species from the family Actinostolidae previously reported in deep areas in Chile have been registered on the Peruvian coastline, but only in the shallow subtidal (Table 1; Alfaro *et al*., 2016; Baldarrago *et al*., 2019). The species *A. achates*, *A. chilensis*, and *P. niveus* have been found in Chile down to 327, 278, and 108 meters respectively (Carlgren 1899, as cited in Häussermann & Försterra, 2009; McMurrich, 1904, as cited in Häussermann & Försterra, 2009; Carlgren 1959, as cited in Häussermann 2004c; Stotz, 1977 as cited in Häussermann & Försterra, 2009). Conversely, in Peru, a unique register of *P. niveus* is available for the locality of Paita (Piura region) without any bathymetric information (Fautin, 2016). In the case of *A. chilensis*, these individuals have been reported inhabiting the same areas as the commercially important sea cucumber *Pattalus mollis* (Selenka 1868) (Alfaro *et al*., 2016). Therefore, a depth range for *A. chilensis* has been obtained as a result of a casual report in the locality of Guañape Island (La Libertad region), yet this information remains unverified in this region, needing further efforts in future research initiatives.

The uneven research effort in the study of sea anemone diversity along the Peruvian coast (Figure 2C-D) may be explained by different factors. The Lima region possesses the highest number of publications referring to sea anemones (Verrill, 1867; C. Paredes & Tarazona, 1980; Tokeshi & Romero, 1995; C. Paredes *et al*., 1999; Häussermann, 2003; Quiroz-Garrido, 2005; Retuerto *et al*., 2007; Firstater *et al*., 2010; Guzmán, 2012; Fautin, 2013; Galán, 2015; Fautin, 2016; Alegre, 2017; Cuya & Escobar, 2017; Tasso *et al*., 2018; Yafac-Piedra & Garcia-Alayo, 2020; A. Ramírez *et al*., 2022; Häussermann, 2004b). One explanation for this is the high research effort produced by several institutions from Lima, where private and public universities, as well as governmental establishments, are involved. Therefore, the Lima region does not only possesses ecological studies (e.g. monitoring reports), as specific research has also targeted sea anemone species to investigate the toxins of their venom (Quiroz-Garrido, 2005) and even its biochemical and biological activity (Retuerto *et al*., 2007; Cuya & Escobar, 2017; Yafac-Piedra & Garcia-Alayo, 2020) to consider it for future pharmaceutical and medical applications. On the other hand, the high number of occurrence registers in the north-central coast (La Libertad and Ancash regions) may be essentially a result of monitoring reports performed as there are determined sites where the intensive culture of Peruvian calico scallop *Argopecten purpuratus* takes place (Uribe *et al*., 2019). Unfortunately, most of the studies consulted do not correspond to a sustained research program of benthic biodiversity but rather episodic or unconnected initiatives. Biodiversity biases associated with the lack of data and information have frequently occurred in the marine environment affecting our ability to analyse specific taxa patterns and dimension ecological responses (Miloslavich *et al*., 2011).

### Taxonomic issues, knowledge gaps and implications for conservation studies

Cnidarian taxonomy in Peru has often been neglected in the past. More specifically, in-depth taxonomy of sea anemones has been overlooked in most studies of marine benthos, paradoxically those of intertidal benthos where this group is a common item. Morphological identification is perhaps the most common limiting factor because of the high intra- and interspecific variability in morphotypes (González-Muñoz *et al*., 2014). But even problematic synonyms have fueled confusion among experts; for instance, *Isoulactis chilensis* (Carlgren, 1959) has been catalogued as a synonym of *O. concinnata*, though they were formerly considered two different species with identical morphological features, colour, habitat, and behaviour (Häussermann, 2003). Additionally, the challenge to differentiate *Actinostola* species even among individuals from the same species has been pointed out as an issue (Häussermann & Försterra, 2009).

Alternative methods such as examining internal morphology through histology (cnidocites) and genetic analyses are needed to perform an accurate taxonomic identification in such cases (Barragán, 2018; Häussermann, 2004a, 2004c). Due to these obstacles, multiple sea anemone species are first registered as *Actinia* sp., like *A. alicemartinae* in Chilean reports from the ’70s (Häussermann & Försterra, 2001). Similar cases hamper the understanding of sea anemones biodiversity in Peru. Indeed, recent monitoring reports registering anemone species by referring to their genus only demonstrate that classifying actiniarians at the species level is still demanding (Pastor *et al*., 2017; De Lucio, 2020).

The identification of sea anemones is essentially based on preserved organisms and old, scattered literature (Häussermann & Försterra, 2009). As an example, the species *A. picta* is still catalogued with an uncertain scientific name (Daly & Fautin, 2021). Issues such as an imprecise original description, doubtful species characters, and biological material (e.g. tentacles) in poor conditions (Verrill, 1867; Andres, 1884) are yet hindering an accurate classification of this species. Future studies in Peru should include detailed *in situ* observations following protocols that encourage observing external characteristics before preservation, as this provides crucial information for the identification (Häussermann, 2004a).

Complementary methods such as metabarcoding may significantly favour our perspective on sea anemone biodiversity. It would give us the potential to identify new species and solve taxonomic doubts for cryptic ones (e.g. *Anthothoe chilensis*, Häussermann & Försterra, 2009), which can affect the quantification of local biodiversity and the assessment of the environment’s status. The relevance of gathering abiotic and biotic data while collecting the individuals has also been highlighted, as this kind of input is still insufficient or unknown for most species inhabiting the Peruvian coast. Accurate information concerning the depth at which organisms have been collected is not unfortunately supplied by the sea anemone literature (Fautin, 2016). Certainly, the actual bathymetric range of most sea anemone species recorded in Peru is still unclear, partially due to the methods used to collect data. The existing studies have mostly used autonomous diving, a method useful for exploring coastal benthic habitats but limited to the diver’s experience, and therefore, with a potential bias on the real bathymetric range.

Comprehensive sampling campaigns targeting sea anemones could positively address the uncertainty existing regarding the habitat range of these specimens, and substantially favouring the determination of spatial patterns of biodiversity and abundance. This is especially crucial when analyzing casual (unverified) reports of sea anemones about which virtually nothing is known. For example, the anemone *Anthopleura mariscali* (Daly & Fautin, 2004) is supposedly inhabiting hard bottoms in the locality of Las Pocitas (Piura region) according to popular websites like iNaturalist. However, these records are the only ones registered for the Peruvian coast and do not have complementary scientific information to corroborate them, revealing further sampling needs and a peer-review analysis. Only after this verification, species such as *A. mariscali* could be officially included in the species list of sea anemones reported in Peru. Similarly, a report of *A. alicemartinae* in the locality of Boca del Río (Tacna region) on the OBIS database requires as well both taxonomic reconfirmation and future *in situ* collection campaigns to validate this observation.

Preliminary reports for species such as *A. radians* indicate a rocky habitat (Puerto Inglés, Moquegua region) as a sporadic distribution (Zuñiga, 2019). Nevertheless, it has been recently indicated this species is abundant in protected and semi-protected areas of rocky intertidal ecosystems on the northern Chilean coast (Spano & Häussermann, 2017), suggesting both a southernmost distribution and an insufficient sampling effort in some Peruvian localities. Also, the possible influence of biological habitats, such as kelp forests, on the biodiversity of sea anemones and their distribution patterns is unknown. For instance, *A. alicemartinae* has been frequently found inhabiting *Macrocystis integrifolia* kelp beds (Villegas *et al*., 2008) and even attached to floating macroalgae in offshore Chilean waters (Thiel & Gutow, 2005) suggesting that this flora is a suitable substrate for anemones but further analyses is needed to elucidate adaptability and habitat range in this group.

On the other hand, particular cases highlight the episodic and/or unconnected research activity on sea anemones but also remarkable opportunities to analyze new data. For example, specimens of *Actinia* sp. and *Paranthus* sp. have been reported as prey items of the green turtle *Chelonia mydas* (Linnaeus, 1758) in both Piura and Ica regions (E. Paredes, 2015; Quiñones *et al*., 2017, 2021a). Patches of *Paranthus* sp. and algae have also been described as part of important feeding areas of *C. mydas* in the Ica region (E. Paredes *et al*., 2020). However, there are cases when some species have not reappeared since their first reports (Verrill, 1867; Andres, 1884; Pax, 1912). Precisely, Pax (1912) suggested considering the sea anemone species *A. picta, P. rubens, Actinothoe gravieri* (*Sagartia gravieri* before), and *Sagartia lessonii* as endemic to Peru because their distribution appears to be restricted to this ecosystem. As the present revision has not hitherto found any register of the occurrence of these species in other countries neither in databases (e.g. GBIF; OBIS) nor in articles, they could be considered as such. Forthcoming research should focus on these species to understand why they have not been spotted again, as we may have ignored these potentially native sea anemone species for decades.

The sea anemone species *A. alicemartinae* may be native to the Peruvian coast. To date, *A. alicemartinae* has only been reported on the Peruvian and Chilean coasts (GBIF; OBIS; Häussermann & Försterra, 2001; Castilla *et al*., 2005; Canales-Aguirre *et al*., 2015; Fautin, 2016; Baldarrago *et al*., 2019; Brante *et al*., 2019; Pinochet *et al*., 2019). Canales-Aguirre *et al*. (2015) suggested that the population present in southern Peru would be the ancestral one that gave rise to the current distribution of *A. alicemartinae* in Chile. Recently, Pinochet *et al*. (2019) concluded this species seems to be invasive to the Chilean coast, coming from the northern Humboldt ecosystem. Until now, the only records of *A. alicemartinae* occurring in Peru are those from the southern coast (Canales-Aguirre *et al*., 2015; Baldarrago *et al*., 2019). Potential studies should be carried out on our coasts to confirm this theory on the origin and distribution of *A. alicemartinae* in the South Pacific. Other sources of bias must be taken into account; namely, the difference in the sampling efforts at regional level (i.e. considering Peru and Chile localities) and its potential influence when analyzing latitudinal distribution patterns and promoting conservation initiatives.

Critically less documented, the potential effects of large-scale ocean-atmosphere fluctuations such as EN are poorly known in Peru, despite being a recurrent event. Recent results suggest diverse ecological responses associated with a significant reduction in the latitudinal distribution range in *P. pluvia* during the 2015 and 2017 EN events, whereas *P. clematis* exhibited an expansion southward (Valqui *et al*., 2021). However, intrinsic biological responses are complex and depend on the degree of adaptability, thermal tolerance, and feeding habits, among other physiological and ecological aspects. Sea temperature variability, for example, is a pervasive driver for survival and can influence the permanence of local populations. Species such as *A. alicemartinae* tend to be more resistant to thermal stress when compared with *Anthothoe chilensis*, which adaptatively responds by increasing its detachment rate to evade higher temperatures (Suárez *et al*., 2020). With both species also reported in Peru, it is likely to observe a similar population response in *A. alicemartinae*, expecting a greater tolerance than *Anthothoe chilensis* amid EN events. In addition, the establishment of *P. clematis* in the lower intertidal zones as a co-dominant species after EN events (1982/83 and 1997/98) has been documented along the northern Chilean coast (Rivadeneira, 1998, as cited in Rivadeneira & Oliva, 2001). The anemone *P. clematis* seems to be a resistant and opportunistic species amidst disturbances in its environment, as there is also evidence of being ubiquitous along the Peruvian coast during EN conditions (Valqui *et al*., 2021). Nevertheless, long-term records of benthic rocky intertidal communities are needed to observe changes attributed to EN events and to have accurate insights concerning benthic responses.

A chronic, shallow and intense OMZ spans most of the Peruvian waters but almost nothing is known regarding the response of sea anemones communities to this stressor. Central Peru holds several representative localities influenced by the presence of OMZ (e.g. San Lorenzo Island) where sea anemones have been observed (see Table 1). Spatially-near findings indicate that local soft-bottom benthic communities are highly adapted to reduced conditions but it is unclear the ecological threshold of these responses over time (Aramayo *et al*., 2021). In fact, future scenarios of climate change for the Peruvian sea highlight that even commercially important benthic species are more threatened than others in this complex marine ecosystem (Ramos *et al*., 2022), consequently producing more uncertainty in other poorly-studied species living in this habitat.

Sea anemone species living in extreme or impacted environments are certainly not rare; Riemann-Zürneck & Gallardo (1990) described a new sea anemone species *Saccactis coliumensis* (now *Oulactis coliumensis*) from samples collected between 40 and 55 m depth in eutrophicated sediments exposed to deoxygenated waters on the central Chilean shelf. As the Eastern South Pacific OMZ comprises the regions of Ecuador, Peru, and Chile (Paulmier *et al*., 2006), it is expected to encounter *O. coliumensis* and perhaps other sea anemone species integrating the Peruvian OMZ benthic fauna.

## CONCLUSIONS

This review is the first produced about this benthic group. Most of the information analyzed and synthesized here is novel to science. It is particularly noteworthy that most of the species reported here do not appear in the usual monitoring reports or the few *ad hoc* studies on this benthic group. Although certainly in recent decades there has been a progressive, albeit slow, growing interest in this group of benthic invertebrates, it is also true that we need to clarify many doubts about species described long ago, and adequately identify the potential biases in this existing data, especially the environmental information which we critically lack.

A significant advance in the study of this group also requires an important effort to analyze its ecological importance and responses concerning the many existing sources of impact, especially large-scale ones such as EN events and regional stressors like OMZ. The increasing frequency of these phenomena in future climate change scenarios is of high possibility, however, studies addressing this panorama are rather insufficient, rare or simply do not exist. Although sea anemones are highly adaptive, more than one of the species reported here may be currently subject to some degree and sort of population pressure, either due to natural causes or anthropic impacts (pollution, habitat loss, etc.) resulting in an increasing uncertainty of the real biodiversity status of this group.

## ACKNOWLEDGEMENTS

This work has been carried out in the research framework of the ECOMARES Project (#B21100601) at UNMSM.

## Notes

### Competing Interest Statement

The authors have declared no competing interest.

## REFERENCES

Alegre, A.R.P. (2017). Cambios en la estructura del megabentos asociado al submareal rocoso en las Islas San Lorenzo y Palomino (Callao, Perú) durante el evento El Niño 1997–98. Universidad Nacional Mayor de San Marcos, Lima, Perú. Retrieved from http://biblioimarpe.imarpe.gob.pe/handle/123456789/3136

Alfaro, S., De Lucio, L., Escudero, L., Atoche, D., Flores, L., Goicochea, C., Campos, M., García, O., and Neira, Ú. (2019). Caracterización de recursos bentónicos en las Islas Guañape (Norte y Sur), La Libertad – 2016. Informe Instituto del Mar del Perú 46, 601–635.

Alfaro, S., Rebaza, V., De Lucio, L., Salcedo, J., and Vásquez, C. (2016). Evaluación de bancos naturales de invertebrados marinos comerciales, Región La Libertad-Perú, 2012. Informe Instituto del Mar del Perú 43, 68–93.

Alfaro, S., Rebaza, V., De Lucio, L., Vásquez, C., and Campos, M. (2017). Caracterización de bancos naturales de invertebrados marinos comerciales y áreas de pesca artesanal. Región La Libertad, Perú. Junio 2014. Informe Instituto del Mar del Perú 44, 105–148.

Andres, A. (1884). Le Attinie (Monografia) (Vol. 1). Leipzig: Verlag von Wilhelm Engelmann.

Aramayo, V., Romero, D., Quipúzcoa, L., Graco, M., Marquina, R., Solís, J., and Velazco, F. (2022). Respuestas del bentos marino frente a El Niño costero 2017 en la plataforma continental de Perú central (Callao, 12°S). Boletin Instituto del Mar del Perú, 36(2), 476–509.

Aramayo, V., Velazco, F., and Solís, J. (in press). Macrobentos de fondo blando somero en la costa sur de Perú (18°10’ – 18°20’ s): características del hábitat, análisis comunitario y distribución espacial de pequeña escala.

Atoche, D. (2017). Biodiversidad de la fauna macrobentónica de las Islas Guañape, Región La Libertad – Perú, 2012-2014 (Master’s thesis). Universidad Nacional de Trujillo, Trujillo, Perú. Retrieved from https://dspace.unitru.edu.pe/bitstream/handle/UNITRU/16183/Atoche%20Suclupe%2c%20Dennis%20Elthon.pdf?sequence=1&isAllowed=y

Baldarrago, D., Aragón, B., Vizcarra, Y., and Tejada, A. (2019). Estructura bentónica en el submareal somero de Punta Coles (Ilo, Moquegua) en el 2017. Informe Instituto del Mar del Perú 46, 578–600.

Barragán, P. (2018). Taxonomía de anémonas de mar (Cnidaria: Anthozoa: Actiniaria) del Pacífico Mexicano y Panameño (Master’s thesis). Universidad Autónoma de Baja California Sur, La Paz, México. Retrieved from http://rep.uabcs.mx:80/handle/23080/254

Barragán, Y., Sanchez, C., and Rodriguez, E. (2019). First inventory of sea anemones (Cnidaria: Actiniaria) from La Paz Bay, southern Gulf of California (Mexico). Zootaxa 4559, 501.

Berrú, P., and Nizama, A. (2016). Ochetostoma baronii (GREEFF, 1879), primer registro para la región Áncash y el Perú. Científica 13, 113–124.

Berrú, P., and Perea de la Matta, Á. (2019). El Niño costero 2017: impacto sobre población de Tagelus dombeii (Lamarck, 1818) y estructura comunitaria del macrobentos en el banco natural de isla Blanca-ENAPU, Perú. Boletín Instituto del Mar del Perú 34, 369–384.

Bertrand, A., Chaigneau, A., Peraltilla, S., Ledesma, J., Graco, M., Monetti, F., and Chavez, F.P. (2011). Oxygen: A Fundamental Property Regulating Pelagic Ecosystem Structure in the Coastal Southeastern Tropical Pacific. PLOS ONE 6, e29558.

Brante, A., Riera, R., and Riquelme, P. (2019). Aggressive interactions between the invasive anemone Anemonia alicemartinae and the native anemone Phymactis papillosa. Aquatic Biology 28, 127–136.

Brusca, R.C. (1980). Common Intertidal Invertebrates of the Gulf of California. Tucson: University of Arizona Press.

Canales-Aguirre, C.B., Quiñones, A., Hernández, C.E., Neill, P.E., and Brante, A. (2015). Population genetics of the invasive cryptogenic anemone, Anemonia alicemartinae, along the southeastern Pacific coast. Journal of Sea Research 102, 1–9.

Carrizo, S. (2014). Cnidocistos de la anémona de mar Bunodactis octoradiata (Carlgren, 1899) (Cnidaria, Actiniaria, Actiniidae): composición, abundancia y biometría. Universidad Nacional de Mar del Plata. Retrieved from https://aquadocs.org/bitstream/handle/1834/14468/Carrizo_2014.pdf?sequence=1&isAllowed=y

Carter, D. (1965). Actinias de Montemar, Valparaíso. Revista de Biología Marina 12, 129–157.

Castilla, J., Uribe, M., Bahamonde, N., Clarke, M., Desqueyroux-Faúndez, R., Kong, I., Moyano, H., Rozbaczylo, N., Santelices, B., Valdovinos, C., and Zavala, P. (2005). Down under the southeastern Pacific: Marine non-indigenous species in Chile. Biological Invasions 7, 213–232.

Chaigneau, A., Dominguez, N., Eldin, G., Vasquez, L., Flores, R., Grados, C., and Echevin, V. (2013). Near-coastal circulation in the Northern Humboldt Current System from shipboard ADCP data. Journal of Geophysical Research: Oceans 118, 5251–5266.

Chintiroglou, Ch., and Koukouras, A. (1992). The feeding habits of three Mediterranean sea anemone species, Anemonia viridis (Forskål), Actinia equina (Linnaeus) and Cereus pedunculatus (Pennant). Helgoländer Meeresuntersuchungen 46, 53–68.

Coronado, K. (2015). Macrozoobentos de la zona intermareal de Punta Uripe, Salaverry 2014. Universidad Nacional de Trujillo, Trujillo, Perú. Retrieved from http://dspace.unitru.edu.pe/handle/UNITRU/4667

Cuya, A., and Escobar, E. (2017). Estudio bioquímico del veneno de la anémona de mar Phymactis papillosa (Actiniidae). Revista Peruana de Biología 24, 303–310.

Daly, M., Brugler, M.R., Cartwright, P., Collins, A.G., Dawson, M.N., Fautin, D.G., France, S.C., Mcfadden, C.S., Opresko, D.M., Rodriguez, E., Romano, S.L., and Stake, J.L. (2007). The phylum Cnidaria: A review of phylogenetic patterns and diversity 300 years after Linnaeus. Zootaxa 1668, 127–182.

Daly, M., and Fautin, D. (2021). World List of Actiniaria. Anactis picta (Lesson, 1830). Accessed through: World Register of Marine Species at: https://www.marinespecies.org/aphia.php?p=taxdetails&id=289406 on 2021-09-24

De Lucio, L. (2020). Bioecología de Chondracanthus chamissoi “yuyo” en las praderas del litoral de Paijan, región La Libertad – Perú, 2015 (Master’s thesis). Universidad Nacional de Trujillo, Trujillo, Perú. Retrieved from http://dspace.unitru.edu.pe/handle/UNITRU/18716

Donayre, S.J. (2021). Influencia de las praderas de macroalgas pardas en la composición de la biodiversidad marina megabentónica en San Juan de Marcona (Master’s thesis). Universidad Nacional José Faustino Sánchez Carrión, Huacho, Perú. Retrieved from http://repositorio.unjfsc.edu.pe/handle/UNJFSC/4527

Duckworth, C.G., Picariello, C.R., Thomason, R.K., Patel, K.S., and Bielmyer-Fraser, G.K. (2017). Responses of the sea anemone, Exaiptasia pallida, to ocean acidification conditions and zinc or nickel exposure. Aquatic Toxicology 182, 120–128.

Excoffon, A.C., Belém, M., Zamponi, M., and Schlenz, E. (1997). The validity of Anthothoe chilensis (Actiniaria, Sagartiidae) and its distribution in Southern Hemisphere. Iheringia, Série Zoologia 82, 107–118.

Fautin, D.G. (2013). Hexacorallians of the world. https://doi.org/10.15468/90drpi accessed via GBIF.org on 2021-03-12

Fautin, D.G. (2016). Catalog to families, genera, and species of orders Actiniaria and Corallimorpharia (Cnidaria: Anthozoa). Zootaxa 4145, 1–449.

Firstater, F.N., Hidalgo, F.J., Lomovasky, B.J., Tarazona, J., Flores, G., and Iribarne, O.O. (2010). Coastal upwelling may overwhelm the effect of sewage discharges in rocky intertidal communities of the Peruvian coast. Marine and Freshwater Research 61, 309–319.

Flores, D.D. (2014). Diversidad de macrozoobentos en Puerto Malabrigo, La Libertad, abril a setiembre 2014. Universidad Nacional de Trujillo, Trujillo, Perú. Retrieved from http://dspace.unitru.edu.pe/handle/UNITRU/12345

Galán, M. (2015). Estructura y composición de la comunidad macrobentónica asociada a la macroalga filamentosa Chaetomorpha crassa en el submareal somero de la Isla San Lorenzo, Callao – Perú. Universidad Nacional Mayor de San Marcos, Lima, Perú. Retrieved from https://cybertesis.unmsm.edu.pe/handle/20.500.12672/9481

Garese, A., Carrizo, S., and Acuña, F.H. (2016). Biometry of sea anemone and corallimorpharian cnidae: statistical distribution and suitable tools for analysis. Zoomorphology 135, 395–404.

Garese, A., Guzmán, H.M., and Acuña, F.H. (2009). Sea Anemones (Cnidaria: Actiniaria and Corallimorpharia) from Panama. Revista de Biología Marina y Oceanografía 44, 791–802.

Gonzáles, A., and Pastor, R. (2017). Comunidades bentónicas de los ecosistemas de fondos blandos y duros en el intermareal y submareal somero. Sitio piloto Punta San Juan. Enero – febrero 2014. Informe Instituto del Mar del Perú 44, 344–370.

González-Muñoz, R., Simões, N., Mascaró, M., Tello-Musi, J.L., Brugler, M.R., and Rodríguez, E. (2014). Morphological and molecular variability of the sea anemone Phymanthus crucifer (Cnidaria, Anthozoa, Actiniaria, Actinoidea). Journal of the Marine Biological Association of the United Kingdom, 95(1), 69–79.

Grajales, A. (2014). Morphological and molecular evolution of sea anemones as revealed by an emerging model organism, Aiptasia (Cnidaria, Actiniaria, Aiptasiidae) (PhD thesis). Richard Gilder Graduate School at the American Museum of Natural History, New York. Retrieved from https://digitallibrary.amnh.org/handle/2246/6717

Grajales, A., and Rodríguez, E. (2014). Morphological revision of the genus Aiptasia and the family Aiptasiidae (Cnidaria, Actiniaria, Metridioidea). Zootaxa 3826, 55–100.

Guzmán, R. (2012). Preferencias alimenticias de tres especies de anémonas (Cnidaria: Anthozoa) del litoral limeño. Museo de Historia Natural “Vera Alleman Haeghebaert” 39–44.

Häussermann, V. (2003). Redescription of Oulactis concinnata (Drayton in Dana, 1846) (Cnidaria: Anthozoa: Actiniidae), an actiniid sea anemone from Chile and Perú with special fighting tentacles; with a preliminary revision of the genera with a “frond-like” marginal ruff. Zoologische Verhandelingen 345, 173–207.

Häussermann, V. (2006). Biodiversity of Chilean sea anemones (Cnidaria: Anthozoa): distribution patterns and zoogeographic implications, including new records for the fjord region. Investigaciones Marinas 34, 23–35.

Häussermann, V. (2004a). Identification and taxonomy of soft-bodied hexacorals exemplified by Chilean sea anemones; including guidelines for sampling, preservation and examination. Journal of the Marine Biological Association of the United Kingdom 84, 931–936.

Häussermann, V. (2004b). Re-description of Phymactis papillosa (Lesson, 1830) and Phymanthea pluvia (Drayton in Dana, 1846) (Cnidaria: Anthozoa), two common actiniid sea anemones from the south east Pacific with a discussion of related genera. Zoologische Mededelingen 78, 345–381.

Häussermann, V. (2004c). The sea anemone genus Actinostola (Verrill 1883): variability and utility of traditional taxonomic features, and a re-description of Actinostola chilensis (McMurrich 1904). Polar Biology 28, 26–38.

Häussermann, V., and Försterra, G. (2001). A new species of sea anemone from Chile, Anemonia alicemartinae n. sp. (Cnidaria: Anthozoa). An invader or an indicator for environmental change in shallow water? Organisms Diversity & Evolution 1, 211–224.

Häussermann, V., and Försterra, G. (2005). Distribution patterns of Chilean shallow-water sea anemones (Cnidaria: Anthozoa: Actiniaria, Corallimorpharia); with a discussion of the taxonomic and zoogeographic relationships between the actinofauna of the South East Pacific, the South West Atlantic and the Antarctic. Scientia Marina 69, 91–102.

Häussermann, V., and Försterra, G. (2009). Fauna marina bentónica de la Patagonia Chilena: guía de identificación ilustrada. Santiago de Chile: Nature in Focus.

Hooker, Y., Ubillús, O., Heaton, J.C., García, O., and García, M. (2011). Evaluación de Objetos de Conservación y Zonificación de Isla Santa, Ancash. Servicio Nacional de Áreas Naturales Protegidas por el Estado (SERNANP – MINAN) 78.

Lancellotti, D.A., and Vasquez, J.A. (2000). Zoogeografía de macroinvertebrados bentónicos de la costa de Chile: contribución para la conservación marina. Revista Chilena de Historia Natural 73, 99–129.

Linton, D., and Warner, G. (2003). Biological indicators in the Caribbean coastal zone and their role in integrated coastal management. Ocean & Coastal Management 46, 261–276.

Miloslavich, P., Klein, E., Díaz, J.M., Hernández, C.E., Bigatti, G., Campos, L., Artigas, F., Castillo, J., Penchaszadeh, P.E., Neill, P.E., Carranza, A., Retana, M.V., Astarloa, J.M.D. de, Lewis, M., Yorio, P., Piriz, M.L., Rodríguez, D., Yoneshigue-Valentin, Y., Gamboa, L., and Martín, A. (2011). Marine Biodiversity in the Atlantic and Pacific Coasts of South America: Knowledge and Gaps. PLOS ONE 6, e14631.

Monroy, A., Lucero, S., Barriga, E., and Quiroz, M. (2020). Bancos naturales de invertebrados marinos en el litoral de Caravelí – Arequipa, 2016. Informe Instituto del Mar del Perú 47, 481–529.

Novoa, J., Hooker, Y., and García, A. (2010). Isla Foca, guía de fauna silvestre (1st ed.). Piura, Perú: Naturaleza y Cultura Internacional – CONCYTEC.

OBIS. (2021). Anemonia alicemartinae Häussermann & Försterra, 2001. Retrieved on 2021-06-04 from https://obis.org/taxon/283318

Ocaña, O., Moro, L., Ortea Rato, J., Espinosa, J., and Caballer Gutiérrez, M. (2007). Guía visual de la biodiversidad marina de Guanahacabibes. I-Anémonas (Anthozoa: Actiniaria, Corallimorpharia, Ceriantharia y Zoanthidea) Visual Guide of the marine biodiversity of Guanahacabibes. I-Anemones (Anthozoa: Actiniaria, Corallimorpharia, Ceriantharia & Zoanthidea). Avicennia 19, 133–142.

Paredes, C., Cardoso, F., and Tarazona, J. (1999). Invertebrados del intermareal rocoso del departamento de Lima, Perú: Una lista comentada de especies. Revista Peruana de Biología 6, 143–151.

Paredes, C., and Tarazona, J. (1980). Las comunidades de mitílidos del mediolitoral rocoso del departamento de Lima. Revista Peruana de Biología 2, 59–72.

Paredes, C., Tarazona, J., Canahuire, E., Romero, L., and Cornejo, O. (1988). Invertebrados Macro-Bentónicos del área de Pisco, Perú. Boletín Extraordinario Instituto del Mar del Perú 121–132.

Paredes, E. (2015). Hábitos alimentarios de la tortuga verde del Pacífico Este Chelonia mydas agassizii (BOUCORT, 1868) en la Bahía de Paracas, Ica, Perú, durante el año 2010. Universidad Nacional Mayor de San Marcos, Perú. Retrieved from https://cybertesis.unmsm.edu.pe/handle/20.500.12672/4369

Paredes, E., Quispe, S., and Quiñones, J. (2020). Tortugas marinas en las islas Ballestas y Chincha, GEF UNDP Perú, 2013. Informe Instituto del Mar del Perú 47, 89–95.

Pastor, R., Gonzáles, A., and Zavalaga, F. (2017). Comunidades bentónicas de los ecosistemas de fondos blandos y duros en el intermareal y submareal somero. Sitio piloto Islas Ballestas. Setiembre-Octubre 2013. Informe Instituto del Mar del Perú 44, 303–331.

Paulmier, A., Ruiz-Pino, D., Garçon, V., and Farías, L. (2006). Maintaining of the Eastern South Pacific Oxygen Minimum Zone (OMZ) off Chile. Geophysical Research Letters 33.

Pax, F. (1912). Les actinies de la côte du Pérou recueillies par le Dr. P. Rivet. In Mission du Service Géographique de l’Armée pour la mesure d’un Arc de Méridien Equatorial en Amérique du Sud sous le contrôle scientifique de l’Académie des Sciences (Vol. 9, pp. 1–28). Paris: Gauthier-Villars, Imprimeur Libraire du Bureau des Longitudes, de l’Ecole Polytechnique.

Pinochet, J., Rivera, R., Neill, P.E., Brante, A., and Hernández, C.E. (2019). Spread of the non-native anemone Anemonia alicemartinae Häussermann & Försterra, 2001 along the Humboldt-current large marine ecosystem: an ecological niche model approach. PeerJ 7, e7156.

Quiñones, J., Quispe, S., and Galindo, O. (2017). Illegal capture and black market trade of sea turtles in Pisco, Peru: the never-ending story. Latin American Journal of Aquatic Research 45, 615–621.

Quiñones, J., Quispe, S., Manrique, M., and Paredes, E. (2021a). Dieta de la tortuga verde del Pacífico este Chelonia mydas agassizii (Boucort, 1868) en el estuario de Virrilá, Sechura-Perú. 2013-2018. Boletin Instituto del Mar del Perú 36, 85–105.

Quiroz-Garrido, Y. (2005). Estudio de las toxinas de la anémona de mar Anthothoe chilensis (Lesson, 1830) (Actiniaria, Sagartiidae). Universidad Nacional Mayor de San Marcos, Lima, Perú. Retrieved from http://cybertesis.unmsm.edu.pe/bitstream/handle/20.500.12672/1404/Quiroz_gy.pdf?sequence=1&isAllowed=y

Ramírez, A., Ganoza Chozo, F., Elliott Rodríguez, W., Gonzales Aranda, P., Silva Silva, G., Fritz Pumachagua, E., and Ramos López, Á. (2019). Bancos naturales de invertebrados y determinación de áreas para maricultura entre Punta Litera y Playa Grande, Región Lima. Informe Instituto del Mar del Perú 46, 162–193.

Ramírez, A., Ganoza, F., Gonzales, R., and Baldeón, A. (2022). Invertebrados bentónicos y peces en bancos naturales entre Ensenada y delta del río Chancay (Provincia Huaral – Región Lima). Informe Instituto del Mar del Perú 49, 108–121.

Ramírez, P., Castañeda, J., De la Cruz, J., Galán, J., and Bances, S. (2017a). Bancos naturales de invertebrados marinos comerciales en el litoral de la Región Lambayeque, Perú. Setiembre 2014. Informe Instituto del Mar del Perú 44, 83–92.

Ramírez, P., De la Cruz, J., and Castro, J. (2020). Biodiversidad marina en Isla Lobos de Tierra, Lambayeque (agosto, 2017). Informe Instituto del Mar del Perú 47, 549–565.

Ramírez, P., De la Cruz, J., and Castro, J. (2019b). Biodiversidad en las Islas Lobos de Afuera, Región Lambayeque. Mayo 2015. Informe Instituto del Mar del Perú 46, 323–340.

Ramírez, P., De la Cruz, J., and Castro, J. (2019c). Biodiversidad marina en Islas Lobos de Afuera, Región Lambayeque, mayo 2016. Informe Instituto del Mar del Perú 46, 426–443.

Ramírez, P., De la Cruz, J., Galán, J., and Castro, J. (2017b). Caracterización de bancos naturales de invertebrados marinos comerciales y áreas de pesca artesanal. Región Lambayeque, Perú. Junio 2014. Informe Instituto del Mar del Perú 44, 63–82.

Ramírez, P., De la Cruz, J., and Torres, D. (2019a). Biodiversidad en la Isla Lobos de Tierra, Región Lambayeque, setiembre 2015. Informe Instituto del Mar del Perú 46, 341–359.

Ramos, J.E., Tam, J., Aramayo, V., Briceño, F.A., Bandin, R., Buitron, B., Cuba, A., Fernandez, E., Flores-Valiente, J., Gomez, E., Jara, H.J., Ñiquen, M., Rujel, J., Salazar, C.M., Sanjinez, M., León, R.I., Nelson, M., Gutiérrez, D., and Pecl, G.T. (2022). Climate vulnerability assessment of key fishery resources in the Northern Humboldt Current System. Scientific Reports 12, 4800.

Retuerto, F., Arbaiza, E., Quiroz-Garrido, Y., Estrada, R., and Zavala, J. (2007). Actividad biológica del veneno de Anthothoe chilensis (Lesson, 1830) (Actiniaria: Sagartiidae). Revista Peruana de Biología 14, 277–282.

Riemann-Zürneck, K., and Gallardo, V.A. (1990). A new species of sea anemone (Saccactis coliumensis n. sp.) living under hypoxic conditions on the central Chilean shelf. Helgoländer Meeresuntersuchungen 44, 445–457.

Rivadeneira, M.M., and Oliva, E. (2001). Patrones asociados a la conducta de desplazamiento local en Phymactis clematis Drayton (Anthozoa: Actiniidae). Revista Chilena de Historia Natural 74, 855–863.

Rodríguez, E., Barbeitos, M.S., Brugler, M.R., Crowley, L.M., Grajales, A., Gusmão, L., Häussermann, V., Reft, A., and Daly, M. (2014). Hidden among Sea Anemones: The First Comprehensive Phylogenetic Reconstruction of the Order Actiniaria (Cnidaria, Anthozoa, Hexacorallia) Reveals a Novel Group of Hexacorals. PLOS ONE 9, e96998.

Ruppert, E.E., and Barnes, B.D. (1996). Zoología de los invertebrados (6th ed.). México: McGraw Hill Interamericana.

Spano, C., and Häussermann, V. (2017). Anthopleura radians, a new species of sea anemone (Cnidaria: Actiniaria: Actiniidae) from northern Chile, with comments on other species of the genus from the South Pacific Ocean. Biodiversity and Natural History 3, 1–11.

Spano, C., Carbajal, P., Ganga, B., Acevedo, C., and Häussermann, V. (2022). Out of the depths: new records of the sea anemone Oulactis coliumensis (Riemann-Zürneck & Gallardo, 1990) in shallow waters from northern Chile and Peru. Journal of the Marine Biological Association of the United Kingdom 1–5.

Stotz, W.B. (1979). Functional morphology and zonation of three species of sea anemones from rocky shores in southern Chile. Marine Biology 50, 181–188.

Tarazona, J., Gutiérrez, D., Paredes, C., and Indacochea, A. (2003). Overview and challenges of Marine Biodiversity Research in Peru. Gayana 67, 206–231.

Tasso, V., El Haddad, M., Assadi, C., Canales, R., Aguirre, L., and Vélez-Zuazo, X. (2018). Macrobenthic fauna from an upwelling coastal area of Peru (Warm Temperate South-eastern Pacific province-Humboldtian ecoregion). Biodiversity Data Journal e28937.

Tejada, A., and Baldarrago, D. (2016). Monitoreo biológico poblacional de Aulacomya atra (Molina, 1782) en el litoral de Moquegua y Tacna, 2014. Informe Instituto del Mar del Perú 43, 46–67.

Thiel, M., and Gutow, L. (2005). The Ecology of Rafting in the Marine Environment. II. The Rafting Organisms and Community. Oceanography and Marine Biology: An Annual Review 279–418.

Tokeshi, M., and Romero, L. (1995). Filling a gap: dynamics of space occupancy on a mussel-dominated subtropical rocky shore. Marine Ecology Progress Series 119, 167–176.

Uribe, R.A., Atoche-Suclupe, D., Paredes, J., and Seclén, J. (2020). Características bioecológicas de la macroalga roja Chondracanthus chamissoi (C. Agardh) Kützing (Rhodophyta, Gigartinaceae) en la zona intermareal del norte del Perú. Boletin Instituto del Mar del Perú 35, 271–293.

Uribe, R.A., Perea, Á., García, V., and Huerto, M. (2019). Biodiversidad marina en el norcentro de la costa de Perú: Un enfoque para la evaluación de planes de manejo 34, 20.

Uribe, R.A., Perea de la Matta, Á., García, V., and Huerto, M. (2019). Biodiversidad marina en el norcentro de la costa de Perú: un enfoque para la evaluación de planes de manejo. Boletín Instituto del Mar del Perú 34, 332–550.

Uribe, R.A., Rubio, J., Carbajal, P., and Berrú, P. (2013). Invertebrados marinos bentónicos del litoral de la Región Áncash, Perú. Boletín Instituto del Mar del Perú 28, 136–293.

Valqui, J., Ibañez-Erquiaga, B., Pacheco, A.S., Wilbur, L., Ochoa, D., Cardich, J., Pérez-Huaranga, M., Salas-Gismondi, R., Pérez, A., Indacochea, A., Avila-Peltroche, J., Ch, M.R., and Carré, M. (2021). Changes in rocky intertidal communities after the 2015 and 2017 El Niño events along the Peruvian coast. Estuarine, Coastal and Shelf Science 250, 107142.

Vassallo-Avalos, A., Acuña, F.H., González-Muñoz, R., and Rivas, G. (2020). New record of Anthopleura radians (Cnidaria: Actiniaria: Actiniidae) from the Mexican Pacific. Latin American Journal of Aquatic Research 48, 869–876.

Verrill, A.E. (1867). Notes on the Radiata in the Museum of Yale College with descriptions of new genera and species. Transactions of the Connecticut Academy of Arts and Sciences 1, 247–351.

Villegas, M.J., Laudien, J., Sielfeld, W., and Arntz, W.E. (2008). Macrocystis integrifolia and Lessonia trabeculata (Laminariales; Phaeophyceae) kelp habitat structures and associated macrobenthic community off northern Chile. Helgoland Marine Research 62, 33–43.

Watson, L.A., Stark, J.S., Johnstone, G.J., Wapstra, E., and Miller, K. (2018). Patterns in the distribution and abundance of sea anemones off Dumont d’Urville Station, Antarctica. Polar Biology 41, 1923–1935.

Yafac-Piedra, N.M., and Garcia-Alayo, F. (2020). Actividad coagulante y de fosfolipasa A2 del veneno de la anémona de mar Phymanthea pluvia (Drayton, 1846) y de la tarántula Grammostola rosea (Walckenaer, 1837). Biotempo 17, 259–267.

Zanabria, U. (2004). Contribución al conocimiento de la zoocenosis de las orillas rocosas. ‘La Ballenita’, Islay – Arequipa 2003. Universidad Nacional de San Agustín, Arequipa, Perú.

Zanabria, U. (2013). Guía de biodiversidad y ecoturismo en la franja marino costera de Arequipa (Organización de Gestión de Destino OGD – Arequipa.).

Zuñiga, J. (2019). Estructura Comunitaria del Macrozoobentos de la Caleta Puerto Inglés, Bahía de Ilo – Moquegua durante el verano del 2017. Universidad Nacional de San Agustín, Arequipa, Perú. Retrieved from http://repositorio.unsa.edu.pe/handle/UNSA/10369

